# Pre-winter fattening and fat loss during wing moult: the annual cycle of fat deposition in captive barnacle geese (*branta leucopsis*)

**DOI:** 10.1101/2020.04.26.062364

**Authors:** Steven J. Portugal, Rona A. McGill, Jonathan A. Green, Patrick J. Butler

## Abstract

Many different physiological changes have been observed in wild waterfowl during the flightless stage of wing moult, including a loss of body mass. Previously we established that captive barnacle geese (*Branta leucopsis*) underwent this characteristic decrease in body mass during their wing moult, even though they had unlimited and unrestricted access to food. In the present study we aimed to determine if this body mass loss during moult comprised mainly a reduction in fat stores, and to ascertain if the captive geese undergo pre-migratory and pre-winter fattening over a similar temporal scale to their wild conspecifics. The non-destructive technique of deuterium oxide isotope dilution was employed to provide repeated measurements of estimated fat deposition from a captive flock of fourteen barnacle geese. Birds were injected with deuterium oxide at 7 distinct intervals for one annual cycle. During the flightless period of the moult, body fat decreased by approximately 40% from the pre-moult value. During late-September and early October, body fat reached its highest point in the annual cycle, both as an absolute value and as a percentage of total body mass. We propose that while the energetic cost of wing moult is not the ultimate cause of fat loss in moulting barnacle geese, the approximate 212 g of fat catabolised during moult would provide sufficient energy to cover the cost of the replacement of the flight feathers, estimated to be 6384 kJ, over an approximate 42 day period. We conclude that the previously recorded increase in metabolism during moult in the geese, led to the use of endogenous fat reserves because the birds reduced rather than increased their food intake rates owing to the increased risk of predation when flightless. We also conclude that captive barnacle geese do undergo pre-winter and pre-migratory fattening, providing further evidence of the innate nature of these fat deposition cycles.

## Introduction

Fat is the primary energy store in birds, and the benefits of fat storage have been widely addressed with respect to the quantity, composition and morphological distribution of fat stored (Blem, 1990; Witter *et al*., 1993). Benefits of fat storage include insulation, mechanical support, protection, buoyancy in aquatic birds and both sexual and social signals. Of most importance, however, is the energy it liberates when metabolised, and it is the most energy-dense substrate in the body (Blem, 1990). Birds utilise fat most during periods of energy deficit, such as during migration, reproduction and moult (Dawson *et al*., 2000).

Geese, like the majority of avian herbivores in the Northern hemisphere, breed in the Arctic and winter in Southern, more temperate areas (Van der Graaf *et al*., 2007; Bonier *et al*., 2007). Being capital breeders, geese have to balance their energy expenditure and food intake throughout the winter months and early spring, to be able to migrate northwards and to breed successfully (van der Graaf *et al*., 2007, Clausen *et al*., 2003). In barnacle geese, *Branta leucopsis*, birds with larger fat stores have been shown to breed more successfully, suggesting fat accumulation during the non-breeding season is central to increasing fitness in this species (Cope, 2003; Portugal *et al*., 2011a; Portugal *et al*., 2011b). Once breeding has been completed, the geese must replace their feathers (i.e. moult) and accumulate sufficient fat reserves for the autumn migration, before climatic conditions on the Arctic tundra deteriorate (Bonier *et al*., 2007). These events all require energy and thus either the utilisation of fat reserves and/or an increase in food intake (hyperphagia).

Waterfowl are part of one of eight orders of birds that undergo a post-breeding simultaneous flight feather moult, rendering them flightless for a period of approximately 15-45 days (Woolfenden, 1967, Hohman *et al*., 1992; Guillemette *et al*., 2012). Studies on wild waterfowl have demonstrated that during this wing moult period, birds lose body mass (Geldenhuys, 1983; Sjöberg, 1986; Van der Jeugd *et al*., 2003; Portugal *et al*., 2009a; Portugal *et al*., 2009b), alter their behaviour (Kahlert *et* al., 1996; Adams *et al*., 2000; ; Portugal *et al*. 2010; Martin and Portugal, 2011) and significantly increase their rate of metabolism (e.g. Guozhen and Hongfa, 1986; Portugal *et al*., 2007; Portugal *et al*., 2019a). Fox and Kahlert (2005) illustrated in greylag geese, *Anser anser*, for example, the majority of this mass loss during moult was a result of a depletion of fat reserves, which they proposed the geese used to meet the shortfall in normal daily energetic requirements (i.e. these stores were expended to supplement exogenous sources of energy in the diet).

Recent work on captive barnacle geese has shown that, despite having constant access to food and exposure to no predators, they respond in a similar way both physiologically and behaviourally to wing moult as their wild conspecifics (Portugal *et al*., 2007). The geese lose approximately 25% of their body mass during the flightless phase, and their rate of resting metabolism increases by 80% compared to non-moult values. Behaviourally, the birds responded to wing moult by significantly increasing their time dedicated to resting, and decreasing the time spent involved in locomotion and foraging (Portugal *et al*., 2007). Looking at the annual cycle of body mass as a whole, of particular note was the increase in body mass in the captive geese during September and early October, coinciding with the pre-migratory and pre-winter fattening period in wild barnacle geese (Portugal *et al*, 2007; Owen *et al*., 1992). Biebach (1996) noted that so far it has been impossible to get captive animals to fatten before the onset of winter, indicating that under the conditions of captivity, factors are missing that, under natural conditions, induce the accumulation of body fat. So far, only year-round body mass has been measured in captive barnacle geese, therefore, it is not certain to what extent periods of mass loss and mass gain are the result of changes in body fat content.

The isotope dilution method is one of the most frequently used non-destructive techniques for determining body composition (Speakman *et al*, 2001). Water is not evenly distributed in body tissues, and proteinaceous tissue contains substantially more water than fat (Speakman *et al*., 2001). Isotope dilution predicts the total body water content by dilution of a stable isotope, and then estimates the total fat content based on the inverse relationship between body fat and body water (Robbins, 1993; Speakman *et* al., 2001; Mata *et al*., 2005). Previously (Portugal 2008), we have demonstrated that, with the appropriate calibration and corrections applied, deuterium oxide isotope dilution can give sufficiently accurate predictions of body fat in captive barnacle geese to study the temporal changes in the ‘population mean’ of a group of birds.

The primary aim of this study was to document year-round total body fat in captive barnacle geese, to establish whether captive birds would follow a similar temporal pattern of fat deposition to that of their wild conspecifics. We also aimed to determine if (a) the geese would deposit fat prior to wing moult, to enable a reduction in foraging and locomotor activities during the vulnerable flightless phase by relying on endogenous fat stores (b) the body mass loss observed in moulting captive geese is primarily a result of a depletion in fat (c) captive geese would deposit fat during September and early October to coincide with pre-migratory and pre-winter fattening in their wild conspecifics.

## Materials and Methods

### Birds and sampling protocol

A captive population of 14 barnacle geese obtained as 3-week old goslings was maintained under natural light in large outdoor aviaries at the University of Birmingham. The goslings were obtained from Bentley Waterfowl Park (Sussex, UK) which has held a self-sustaining captive population of this species since 1982. The geese were fed with a 50-50 diet (Lilico, Surrey, UK) of mixed poultry corn (4% fat, 12% protein and 71% carbohydrate) and poultry growers pellets (3% fat, 16% protein and 61% carbohydrate), and water was available *ad libitum*.

To provide estimates of body fat content from periods of relatively stable body mass and periods of body mass change, sampling periods were selected based on year round body mass data collected from the same captive flock of barnacle geese the previous year (Portugal *et al*., 2007). Birds were sampled in February (n = 7), March (n = 14), June (n = 11), July (n = 5), August (n = 5), October (n = 7) and December (n = 5), of 2005. Birds were sampled at approximately the same time of day (09:00 – 12:00 GMT) to control for any natural daily rhythms and minor changes in body mass and fat content (Speakman *et al*., 2001). Sample numbers vary because of samples drying out while in capillaries and storage.

### Isotope dosage and administration

For each goose, food was withheld for 7 hours prior to deuterium oxide (D_2_O, Sigma-Aldrich, 99.98%) administration and the sampling of blood for D_2_O analysis.

D_2_O dosage was calculated using equation 12.1 from Speakman (1997);

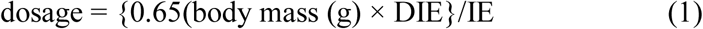

where the constant, 0.65, is the approximate proportion of the body comprised of water, DIE is the desirable initial enrichment (ppm) and IE is the injectate enrichment (ppm). Birds were weighed to the nearest 5 g. Blood samples were taken from either the brachial or intertarsal vein for determination of background enrichment of D_2_O, immediately prior to D_2_O administration. A disposable 1 ml syringe (Terumo) fitted with a 24-gauge stainless steel hypodermic needle was used for administration of the D_2_O, via an intra-peritoneal injection into the lower abdomen. D_2_O was injected slowly and the needle left in for five seconds, to avoid any injectate seeping out (Speakman *et al*., 2001). The actual amount of D_2_O injected was accurately determined by weighing the syringe to the nearest 0.0001 g before and after injection. Subsequent blood samples were collected 90 min after administration of the D_2_O, based on pilot data showing that 90 min was an appropriate length of time for the isotope to equilibrate (Portugal 2008). Blood samples were drawn up in 50 μL non-heperinised glass capillaries (Vitrex, Cambridge, UK) and immediately shaken to the centre of the capillary, which was flame sealed with a butane gas torch (Radio Spares, Corby, UK) and then wax sealed. Capillaries were then stored in an air-tight container.

### Calculation of ^2^H dilution space

Isotope dilution space was calculated by the plateau method (Halliday and Miller, 1977), using equation 4 of Speakman *et al*. (2001);

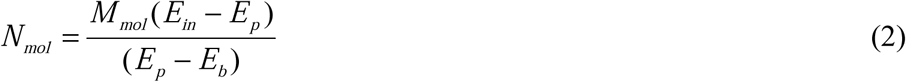

where N_mol_ is the molar quantity of water present in the body, E_in_ is the enrichment of the material introduced into the animal, M_mol_ the molecular weight of the D_2_O, E_b_ is the background enrichment of this material in the animal, and E_p_ the enrichment measured after the ‘dispersal’ process is completed.

Assuming the constancy of the water content of the lean body mass (LBM; Pace and Rathbun, 1945), estimates of body composition could be obtained as follows:

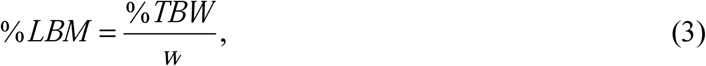

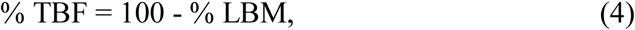

where total body fat (TBF), total body water (TBW) and LBM were expressed as a percentage of fresh body mass. Here, *w* is the mean fractional water content of LBM for nine captive geese that were sampled destructively (0.731 ± 0.013, Portugal 2008).

TBW determined via D_2_O dilution (hereafter referred to as TBW D_2_O) was corrected (e.g. Mata *et al*., 2005) using the relationship between TBW D_2_O and TBW obtained through desiccation (hereafter referred to as TBW DES), before TBF was determined. This correction factor was achieved through calibrating the two (Portugal 2008). The relationship between TBW DES and TBW D_2_O was significantly different between the two sexes, and thus TBW D_2_O for each sex was corrected as follows:

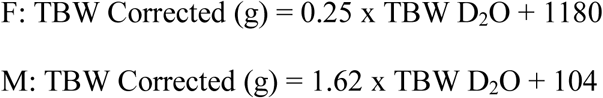

### Isotope analysis

D_2_O enrichment was measured using a chromium reduction furnace interfaced with a dual-inlet isotope-ratio mass spectrometer (Donnelly *et al*., 2001), at the Scottish Universities Environmental Research Centre (SUERC), East Kilbride. Water was extracted from the blood samples by vacuum lyophilisation. Laboratory water standards were also prepared in the same way to correct for day-to-day variations in the performance of the vacuum line. Samples and standards were collected in 50 μL non-heperinised glass capillaries, shaken to the centre of the capillary, and flame sealed with an acetylene torch. Capillaries were broken and the extracted water was then injected in to the mass spectrometer (VG Optima) (after Donnelly *et al*., 2001). Each batch of samples was analysed with triplicates of three laboratory standards, waters of known isotope composition (calibrated relative to SLAP, SMOW and GISP international water standards). These lab standards perform a dual purpose to both correct the isotope ratio of the sample relative to the international standards and to correct for daily mass spectrometer drift, and day-to-day variations in the performance of the mass spectrometer. Isotopically characterised H_2_ gas was used in the reference channel. All isotope enrichments were measured in delta (per mil) relative to the reference gas and those values are in turn calibrated by the laboratory standards, and converted to ppm using established ratios for the reference materials. Measures of isotope enrichment were based on independent analysis of at least two sub-samples of the water extracted from the blood samples.

### Statistical analysis

Since TBW D_2_O was corrected using the calibration relationship between TBW D_2_O and TBW DES, estimates of meant TBW and hence TBF are presented ± the standard error of the estimate (SEE, Zar, 1984). Estimates of fat content were compared using Woolf’s test for differences with Bonferonni corrected proximate normal test (*Z-*test) post-hoc comparisons (e.g. Green *et al*., 2002).

## Results

Body mass changed significantly throughout the annual cycle (Fig. 1a, ANOVA, *P* < 0.001). Estimated total body fat content of the captive barnacle geese changed significantly between the seven sampling sessions (Fig. 1b, Woolf’s test for equality, χ ^2^ = 68.8, *P* < 0.0001). Further *Z*-tests, with Bonferonni corrections, between the seven sampling sessions showed that total body fat content during wing moult in August (129.9 ± 22.4 g) was significantly lower than each of the other six sampling sessions. Between early July and mid-August, the geese lost approximately 212 g, or 40%, of body fat over a 5-week period, an average daily fat loss of 6 g. Fat content was at its highest in the autumn, where estimated fat content peaked at 352.1 ± 20.9 g. Consequently, over the 42 days between the August and October sampling sessions, the geese gained, on average, approximately 5.2 g per day. Body fat decreased steadily during the winter months, with total body fat values of 234.1 ± 23.4 g and 189.5 ±24.9 g being recorded for December and February respectively (Fig. 1b). Fat content in February represented a 53% decrease on body fat at the start of the non-breeding season in October, and fat content was significantly lower in February than both July and October. Between February and March (280.9 ± 17.1 g), body fat increased by approximately 91 g, before reaching a peak just prior to wing moult in July (342.5 ± 36.6 g). Fat as a percentage of body mass followed a similar pattern to that of actual fat mass (Fig. 1c), with body fat percentage ranging from 20.2% in October to just 8.3% in August.

**Figure 1:**
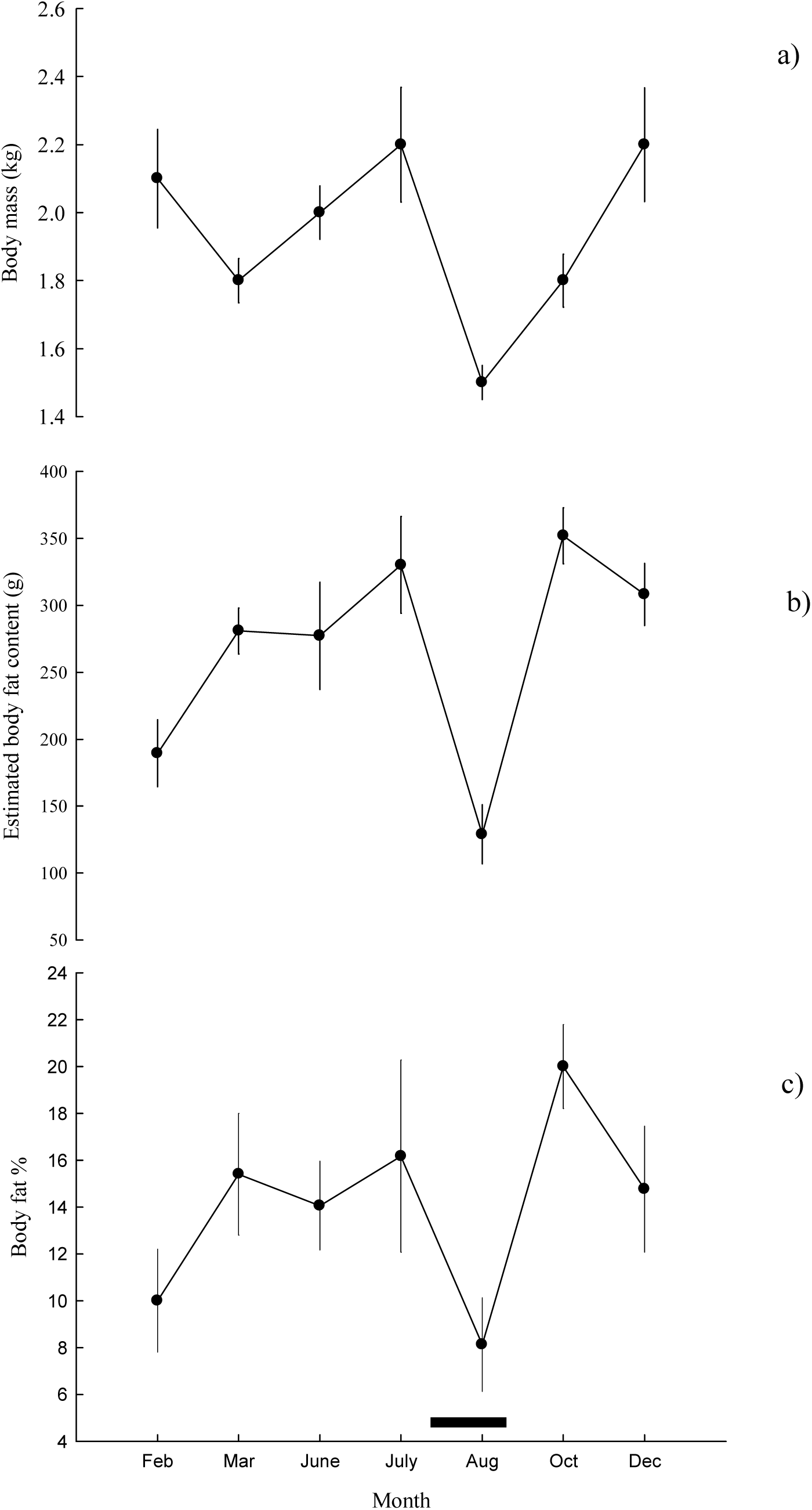
(a) Year-round body mass (± SEM), (b) fat content (± SEE) and (c) fat content as a % (± confidence intervals) of total body mass, from captive barnacle geese. Fat content changed significantly throughout the annual cycle (Woolf’s test for equality, χ^2^ = 68.8, *P* < 0.0001). Black bar represents approximate wing moult period.

## Discussion

### Fat loss during wing moult

Despite constant access to food, the captive barnacle geese lost approximately 212 g of body fat during the wing moult period, equivalent to 40% of their fat reserves in comparison to pre-moulting levels in July. By the end of wing moult in mid-August, body fat only constituted on average 8% of the bird’s total body mass, compared to 16% and 22% pre-and post-wing moult. Fox and Kahlert (2005) recorded a loss of 268 g of fat on average in wild moulting greylag geese, however, no changes in fat content during moult have been identified in mallard *Anas platyrhynchos*, great-crested grebe *Podiceps crisatus*, common scoter *Melanitta nigra* or lesser scaup *Aythya affinis* (Young and Boag, 1982; Piersma, 1988; Fox *et al*., 2008; Austin and Fredrickson, 1987).

Depletion of fat stores during moult in wild birds, like body mass loss, has been interpreted to be a result of nutritional stress caused by the regrowth of the feathers (e.g. Hanson, 1962), a response to the increased risk of predation brought about by being flightless (e.g. Panek and Majewski, 1990) and an adaptation to regain flight quicker on partially regrown flight feathers (e.g. Owen and Ogilvie, 1979). Previously, we concluded that in barnacle geese at least, mass loss (and thus fat loss in this instance) was a consequence of the 80% increase in metabolism during moult, coupled with the reduction in time spent foraging, causing a decrease in body mass (Portugal *et al*., 2007). Therefore, the proximate cause of fat loss is likely to be the interaction of the decrease in food intake and the increase in metabolic rate as a result of feather synthesis. However, the ultimate cause of fat loss in wild birds is the increased risk of predation brought about by the bird’s flightless state, restricting the ability to forage, meaning the geese are not able to increase their food intake to counteract the observed increase in metabolism. As a result, they must rely on their endogenous fat reserves during the flightless phase of moult.

Further evidence that this change in behaviour and subsequent loss of body fat is a product of flightlessness and the increased risk of predation, stems from the increase in body mass and body fat in July just prior to wing moult. At this stage, the captive geese were undergoing their annual body moult, which stretched over a period of approximately 90 days between July and September. Despite regrowing heavier body feathers at this time (e.g. Panek and Majewski, 1990), the geese were gaining body fat, suggesting the cost of regrowing feather alone is not sufficient to result in a loss of body fat.

Although most of species of duck studied do not lose significant amounts of fat during wing moult, certain species of geese have been shown to (Hohman *et al*. 1992). Therefore, it seems likely that there is a link between foraging style, food type and fat loss during moult, and choice of moult site. Many species of duck can continue foraging throughout the duration of moult, as their food source is obtained either from the surface, or under the water, where the birds are relatively safe. Moulting barnacle geese in the Arctic rely on vegetation that is not found in water, and can often be some distance from a water body (Owen and Ogilvie, 1979). Therefore, for the geese, resting on or beside water for safety is frequently mutually exclusive with productive foraging. The fat, and hence and body mass, increases just prior to moult are likely to serve a dual function of enabling the geese to reduce foraging when flightless and rely on endogenous reserves, while increasing the rate of feather synthesis during moult as more reserves are available for this, thus potentially shortening the flightless period – this has been suggested as the possible cause of mass increase prior to wing moult in some duck species (e.g. van Weetering and Cooke, 1999).

Potentially, geese may be able to build large fat stores prior to moult because of their larger size. For example, in coots and grebes, structurally larger individuals can survive longer without food, while being able to deposit larger fat stores (Biebach, 1996). It may be that ducks, being smaller, are unable to deposit sufficient fat stores, and therefore must continue foraging while flightless, albeit at a reduced level (e.g. Adams *et al*., 2001).

Fox and Kahlert (2005) described body mass and fat loss in moulting greylag geese as a representation of the nutritional or energetic “stress” of wing moult, and proposed it to be a further example of avian phenotypic plasticity. This trait enables the geese to meet a nutritional or energetic shortfall from endogenous reserves, in a way that may not necessarily be related to direct fitness costs. Therefore, the barnacle geese show adaptive fat gain prior to wing moult, and the accumulation of fat stores will enable the geese to exploit habitats that are predator free, but which perhaps do not provide the required levels of exogenous nutrient or energy to sustain them through moult.

### Does fat catabolism cover the cost of moult?

The captive geese lost approximately 200 g of fat during late July and August. Although it is unlikely that the energetic cost of wing moult is the ultimate cause of body mass loss in moulting waterfowl, the increase in metabolism coupled with a decrease in foraging work together to cause a loss in body mass, and in this instance, a depletion of fat stores. Using simple calculations it is possible to calculate what the cost of wing moult in the geese would be, and to see if the fat catabolised would provide sufficient enegry to ‘cover the cost’ of the regrowth of the flight feathers.

Plumage mass of birds can be roughly approximated by the equation, plumage mass = 0.09*W*^0.95^ (Turcek, 1966; Lindström *et al*., 1993; Murphy, 1996), and mass of plumage is typically around 6% of the total body mass (Turcek, 1966). As demonstrated in Portugal *et al*. (2007) captive barnacle geese have a large annual variation in body mass, therefore, the average mass for all geese for the month of April 2005 (when body mass is stable, average of 1.8 kg) was used to determine approximate average plumage mass, and the value calculated was 112.37 g (6.1% of total body mass). With the associated mass of the feather sheath included (Murphy, 1996), the total plumage mass for a captive barnacle goose is 134.4 g (for comparison, Lindström *et al*., 1993, cited a plumage mass of 160.6 g for the larger greylag goose). Flight feathers typically account for about one-quarter to one-third of the total plumage mass (e.g. Newton, 1966). The midpoint of these two estimates would provide a value of 36.1g for flight feathers alone for the captive barnacle geese, or 43.2 g including sheaths.

Using equation 3 of Lindström *et al*. (1993) it is possible to calculate the rate of feather production (*C*_f_) if mass specific metabolic rate (kJ g^-1^ d^-1^) of the bird is known. Coupled with plumage mass, it is then possible to calculate the energetic cost of replacing all feathers, and thus the cost of moult. For the captive geese in the present study, rate of feather production was calculated to be 147.8 kJ (g dry feather) ^-1^. C_*f*_ typically decreases with increasing body size, for example values reported in Lindström *et al*. (1993) demonstrated 26.6 kJ (g dry feather)^-1^ for the ostrich, *Struthio camelus*, and 780 kJ (g dry feather)^-1^ for the goldcrest, *Regulus regulus*.

With the mass of flight feathers and the rate of C_*f*_ known, the overall cost of producing the flight feathers can be estimated to be 6385 kJ, or 152 kJ d^-1^ for the approximate 42 day duration of wing moult (assuming feather growth is constant throughout a 24-hour period). If the energy density of fat is 39 kJ g^-1^, 163 g of fat on average would be required to cover the ‘cost’ of re-growing the flight feathers. It is likely, therefore, that most of the 212 g of fat catabolised during the wing moult period was used to grow the new flight feathers. As foraging was greatly reduced (Portugal *et al*., 2007) during wing moult, it is likely the remaining 35 g or so of fat will contribute towards the shortfall in food intake.

### Pre-winter fattening

Body fat made up the highest percentage of total body mass (21%) in late September – early October, before decreasing in December and then reaching the lowest point of the year in February. Biebach (1996) commented that it had not been possible to get captive animals to fatten prior to winter, however, the high percentage of body fat in autumn in the captive geese suggests that this was the case in the present study. If body fat percentage following wing moult returned to a similar level to that of late spring and early summer (e.g. 15%) it could be considered that body fat was returning to ‘normal’ fat levels after fat loss during wing moult. However, as post moult fat was higher both in percentage and absolute terms, it suggests that there is an element of pre-winter fattening taking place. This provides further evidence that captive geese undergo the same physiological changes as their wild conspecifics.

In the Svalbard population of wild barnacle geese, overall fat stores increased and then varied throughout the non-breeding season, with the rate of increase in fat stores being most rapid at the start and end of the non-breeding period (Cope, 2003). Owen *et al*. (1992) found that fat accumulated in wild barnacle geese during the autumn, followed by a significant decline throughout winter and an increase late spring. Phillips *et al*. (2003) noted that the geese depleted the most profitable feeding areas by around early December, after which they switched to less profitable feeding areas, and subsequently lost body fat. During the non-breeding season, peaks in abdominal profile indexes (Owen, 1981) occurred in October and November, with the lowest point coming in February.

The captive geese followed a similar pattern to the wild geese. This pattern suggests that fat levels and control of fat follows an endogenous cycle, being only slightly modified by environmental constraints. Owen *et al*. (1992) suggested that birds might lay down reserves in preparation for lean periods, and then only maintain sufficient reserves to guarantee against predictable adversity. It is possible therefore, that once the geese have arrived in Scotland and recovered their body fat reserves from their autumn migration, a loss of unnecessary reserves may be advantageous to reduce the risk of predation, as heavier birds are probably less agile (Portugal *et al*., 2011c).

The one highly unpredictable event for the geese will be the conditions on arrival on their wintering grounds in Scotland following migration. During spring migration the geese take roughly 4-5 weeks to reach their breeding grounds, essentially following the new growth of vegetation northwards (Cope, 2003; Portugal *et al*., 2019b). Fattening for the spring migration is also a two-step process, first on the Solway and then on Helgeland, a major staging area in Norway (Black *et al*., 1991). Here, fat reserves expended during the first part of the migration are at least partially replenished. In autumn however, geese migrate from Svalbard to Scotland in far shorter time, with some birds capable of achieving this is less than one week (Butler *et al*., 1998). Unlike the situation in spring, geese migrating in autumn do not know what conditions will be like on arrival at their destination. Therefore, large fat reserves are essential not only for migration but as an insurance for arrival at the wintering grounds, should conditions be adverse. Butler and Woakes (2002) demonstrated that wild barnacle geese undergo a period of seasonal hypothermia (or anapyrexia) in autumn thought to aid in fat deposition, not only prior to migration but for recovery of fat reserves once the wintering grounds are reached, again stressing the importance of pre-migratory and pre-winter fattening in this species.

In the present study we have demonstrated that captive barnacle geese do deposit fat reserves prior to wing moult, and that subsequently the majority of mass loss during the flightless phase of wing feather replacement is primarily a depletion of fat. In October, fat reserves in the geese increased significantly, suggesting that the birds were depositing fat, before the onset of winter.

## Acknowledgements

Isotope work was possible thanks to a grant from NERC to use facilities at the SUERC. We would like to thank Jason Newton, Alison MacDonald and Terry Donnelly (SUERC) for their assistance with water extraction and equipment. We are grateful to Alan Gardner, Phil Archer, Ben Heanue and Pete Jones for looking after the geese. Thanks also to Craig White, David Gardiner, Alex Kabat, and Claire Tyler for holding birds during blood sampling sessions.

